# Rethinking Statistical Learning as a Dynamic Stochastic Process, from The Motor Systems Perspective

**DOI:** 10.1101/2022.01.18.476831

**Authors:** Anna Vaskevich, Elizabeth B Torres

## Abstract

The brain integrates streams of sensory input and builds accurate predictions, while arriving at stable percepts under disparate time scales. This stochastic process bears different dynamics for different people, yet statistical learning (SL) currently averages out, as noise, individual fluctuations in data streams registered from the brain as the person learns. We here adopt the motor systems perspective to reframe SL. Specifically, we rethink this problem using the demands that the person’s brain faces to predict, and control variations in biorhythmic activity akin to those present in bodily motions. This new approach harnesses gross data as the important signals, to reassess how individuals learn predictive information in stable and unstable environments. We find two types of learners: narrow-variance learners, who retain explicit knowledge of the regularity embedded in the stimuli -the goal. They seem to use an *error-correction* strategy steadily present in both stable and unstable cases. In contrast, broad-variance learners emerge only in the unstable environment. They undergo an initial period of memoryless learning characterized by a gamma process that starts out exponentially distributed but converges to Gaussian. We coin this mode *exploratory*, preceding the more general error-correction mode characterized by skewed-to-symmetric distributions and higher signal content from the start. Our work demonstrates that statistical learning is a highly dynamic and stochastic process, unfolding at different time scales, and evolving distinct learning strategies on demand.

## Introduction

At the start of life, human babies gradually become aware of their bodies in motion and as they understand it, they come to own the consequences of impending movements that make up all their purposeful actions. Neonates explore their surroundings as they expand their limbs with antigravity motions and eventually learn to reach out to their immediate space in a well-controlled, intended manner. This type of highly dynamic, spontaneous, exploratory learning that is at first driven by surprise and curiosity, has no initial goal or desired target. It is merely a wondering process, “*what happens if I do this?*”, perhaps a guess, “*if I do this, then this (consequence) will ensue, otherwise, this other (consequence) will happen…*” Its endogenous and dynamic nature in early life may scaffold how we start to learn in general, before realizing certain regularities leading to desired goals and conducive of a predictive code.

Research about learning, whether in the perceptual, the motor, or the cognitive domain, is nevertheless primarily based on error-correction schemas ^1-3^. These schemas are aimed at reducing the difference between a desired configuration or goal to be learned, and the current learning state ^3^. Such goals tend to be exogenous in nature, but implicit in them is a regularity that the system must find. Somehow the spontaneous self-discovery process that we relied on as babies, *i*.*e*., to learn about sensing our body in the world and sensing the world in our body, tends to fade away from our behavioral research. Indeed, curious exploration seldom enters our experimental paradigms in explicit ways ^4^. Some animal models of exploratory behavior ^5^ have nevertheless been successfully extended to characterize exploration in human infants as excursions that separate segments of movements’ progression from lingering episodes ^6^. This recent research suggests behavioral homology across species and prompted us to hypothesize that at a finer temporal learning scale, a wondering, exploratory code may hide embedded in the fluctuations of our performance. We tend to average out such fluctuations as superfluous noise. Certainly, when favoring *a priori* imposed theoretical means under assumptions of normality and stationarity in the data registered during the learning process, we miss the opportunity to know what possible information lies in that gross data.

Notably, the cognitive processes known as implicit or statistical learning (SL) describe the ability of the brain to extract (largely beneath awareness) patterns based on regularities from the environment over time ^2,3,7^. Such capacity has long been known to support a wide range of basic human tasks such as discrimination, categorization, and segmentation of continuous information ^8^, thus leading the way in understanding language acquisition ^9,10^. and predictive aspects of social interactions ^11,12^. Previous research has consistently shown that regardless of the nature of the embedded regularity (motor, perceptual or both), statistical learning involves motor control systems, so that when participants are required to respond, the presence of predictive information modulates both response preparation and response execution processes ^13-15^. Yet, very little work addresses the stochastic motor signatures scaffolding statistical learning during *e*.*g*., motor decisions to communicate a preferred stimulus ^16,17^. Prior cross-sectional research points at the dynamically changing stochastic signatures of learning that unfolds at different time scales of maturation. Particularly interesting is that such maturation transitions do not take place in the intended movements of autistics ^16^. They occur in the minute moment-to-moment fluctuations of their spontaneous, exploratory hand movements in search of a surprising audio-visual event ^18-20^. Such movements do not pursue a specific goal and bear similar signatures in autism to those experienced by neurotypical participants ^20^. This prior research inspired us here to reevaluate statistical learning from the standpoint of sensory-motor systems and the different movement classes that the brain recruits on demand, according to levels of intent and deliberateness ^19,21^. We reasoned that the motor percept that emerges from the sensations of our own endogenously generated movements could serve to support the type of statistical learning that other perceptual systems relying on exogenous sensory inputs would experience to gain behavioral control. After all, behavior is neural control of movement at the central and the peripheral levels of the nervous systems.

We here propose to reframe the statistical learning problem by recent advances in developmental research of neuromotor control. To that end, we rely on methods developed at the Torres lab ^16,19,21-23^, and track the dynamic signature of the learning process, continuously evaluating an electroencephalographic (EEG) signal recorded while participants perform in a learning task that contained predictive information. We specifically consider multiple time scales and different levels of stochastic control present in the motor code that scaffolds deliberate goal-directed actions and goal-less exploratory motions. The latter, spontaneously evoked by unexpected events. For any given learning task where these modes coexist, the processes involved may range from local processes at sub-second time scale, to global consolidation processes at the time scale of hours, days and even years (*e*.*g*., as in Figure 1A specific to this experimental paradigm). Within the statistical learning domain, focusing on the dynamics of the learning process itself, with the specific consideration of multiple time scales, has been recently suggested as the next necessary step in statistical learning research ^3,4,7^. That is, to understand the neural substrates underlying behavior it is necessary to view it, and to measure it, as a continuous process, evaluating learning trajectories of its stochastic variations and learning stability. However, so far, these suggestions have not matured into meaningful research, largely due to limitations of the standard analytical techniques.

**Figure 1.**
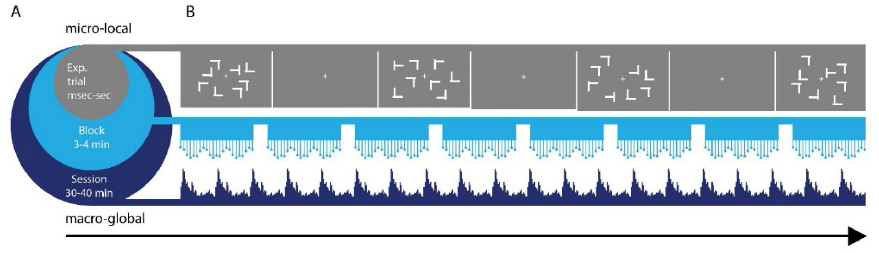
Panel A. Statistical learning paradigm. Different time scales of learning are accompanied by different types of learning supporting each level. From a micro level at sub-second time scales, to a macro level at the scale of 40 minutes (in this experiment), different levels of granularity in the data afford different levels of precision to describe learning phenomena. Averaging out fluctuations in the system’s responses may eliminate gross data containing important information on learning mechanisms. These may be varying from trial to trial and from block to block at each level. Panel B. Visual search task: the target was a letter T rotated either left or right that appeared among rotated Ls (distractors). Across trials, the spatial configurations of target and distractors (*i*.*e*., layouts) could repeat (correlated group), be generated randomly (random group) or repeat on half of the trials (mixed group). EEG signal was recorded while participants searched for the target and pressed the corresponding response key as fast as possible.

A common problem that we encounter in data analyses in general, is that continuous data streams registering the evolution of our brain responses to the learning process, are often summarized using statistical models that make inferences about the learning phenomena under assumptions of linearity, normality and stationarity in data that is inherently nonlinear, non-normally distributed and non-stationary. In the field of statistical learning exclusive reliance on error-correction learning mode, converging to a predictive code, misses the opportunity to uncover a possible exploratory phase of the learning, reminiscent of the type of learning experienced earlier in life, during our infancy, and present thereafter in the motor code of natural behaviors. The error-correction learning mode precludes self-initiate surprise, *i*.*e*., unexpected events possibly leading to wondering mental curiosity. In the motor code, these are rather conducive of signatures of contextual variability unique to each person ^21,24^. Such mental processes are embodied and situated ^25^, *i*.*e*., grounded on how we once learned to behave as infants, through our dynamic bodies in constant (endogenously generated) exploratory motion, reactive to exogenous stimuli.

We take in this work a radically different approach to the types of analyses of data derived from a perceptual learning paradigm currently employed in statistical learning ^3,4,7^. While we leverage the precise time stamping of the events in the data acquisition system and the use of stable and unstable implicit-learning environments ^15^, we convert the continuous EEG data stream to spike trains (coined henceforth micro-movement spikes, MMS). These MMS standardize the signal and scale out anatomical disparities across participants, to leverage the playfield and appropriately compare performance across the group. This approach to data analysis is akin to converting analogue to digital signals and continuously tracking the moment-to-moment shifts in stochastic signatures of the data that are empirically estimated, rather than theoretically assumed. We let the data speak and reveal to us the primordial way of curious, exploratory learning, preceding the discovery of regularities conducive of an error-correction mode. We reframe statistical learning from the point of view of a developing, nascent motor system.

## Results

We use the visual search, a common perceptual task (Figure 1B and see methods for details) where participants (70 in total) search for a ‘T’ (target) embedded among heterogeneously rotated ‘L’ distractors. The participants were randomly distributed among 3 groups where the degree of regularity in the task varied along a gradient. At one extreme the participants searched for the target within a predictable environment where predefined spatial configurations of target and distractors (layouts) were repeated from trial to trial (*i*.*e*., consistent mapping). In this group (coined correlated), the embedded regularity facilitated performance, so that participants reached the fastest reaction times. At the other extreme, in the random group, participants experienced the least amount of regularity, as from trial to trial, the layouts of the display were generated randomly. Here participants reached slower reaction time, although significant perceptual learning still occurred. The results from the analyses of the behavior and averaged potentials were previously reported ^15^. Here we focus on a radically different approach to the full EEG data.

The two above-described groups represent two extreme cases that are artificially constructed for laboratory purposes. The third group, coined mixed, consisted of a more realistic global environment, with both consistent and random conditions intermixed (classic contextual cueing paradigm ^26^. Although the mixed group contained a potentially beneficial regularity on half of the trials, participants reached the slowest performance ^15^. This pattern of results highlights the crucial issue of validity: when the regularity is valid, applying this statistical information results in facilitation to both the search and response processes. However, when the regularity is mixed with random trials, thus appearing within a relatively unreliable and unstable environment, a global interference effect emerges, so that the reliance on all prior information is attenuated (for a detailed account of the origins of this interference effect see ^15,27^.

This task is ideal for the present purposes as it enables to examine stochastic variations in learning between environments that differ in the reliability of the statistical information, while the perceptual input remains the same. As participants performed the task of identifying the ‘T’ among the ‘L’s, the EEG patterns were continuously registered with breaks to rest, avoiding fatigue. Registered at 256Hz, we converted the fluctuations in the EEG amplitude (peaks *μV*) and inter-peak-interval timing (*ms*) to (unitless, standardized) MMS trains (see methods.) We then tracked their stochastic shifts in amplitude and timing, at the local and at the global time scale levels. Locally, we considered trial by trial across 8 blocks of learning (defined by the breaks.) This was on the order of sub-seconds to minutes. Globally, we pooled across all trials in each block and unfolded the data block by block. This was on the order of approximately 40 min of the full experimental session (Figure 1A).

**Local level** of analyses revealed two fundamentally different types of learners in the mixed group. Figure 2A shows the mixed group split for targets oriented to the right and to the left, with the response corresponding to the orientation (right hand for oriented right target). Figures 2B and 2C show the results for the correlated and random groups distinguishing the left and right targets with different colors. Panels in Figure 2D show the frequency histograms obtained by pooling the variance across all subjects for both target types in each group. The mixed group is indeed significantly non-unimodal, according to the Hartigan dip test of unimodality, *p<0*.*01* ^28^. These PDFs significantly differ from those in the random and correlated groups, according to the Kolmogorov Smirnov test for two empirical distributions *(p<0*.*01)*.

**Figure 2.**
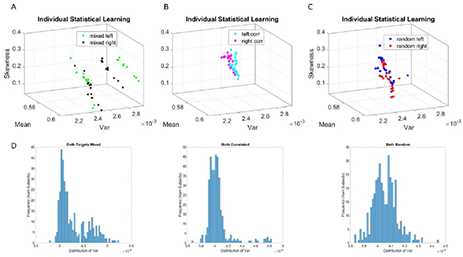
Local learning evolution captures two classes of learners in the unstable environment (*i*.*e*., mixed group). Empirically estimated Gamma moments span a parameter space whereby each participant represents a point by the moments of the probability distribution. The coordinates are the mean (*x-axis*), the variance (*y-axis*), the skewness (*z-axis*). The color represents the target orientation (left or right.) (A) Mixed case *(i*.*e*., group) whereby trials intermix random and correlated conditions, spanning a relatively unstable learning environment. In this group two self-emerging distinct subgroups of participants were distinguishable by target type. (B) Correlated group, for which layouts are consistent from trial to trial, spanning a stable learning environment. (C) Random group, for which layouts are generated randomly from trial to trial, spanning a stable learning environment where no regularity is present. (D) Corresponding frequency histograms of the distribution of the variance across trials, target types and participants.

Figure 3 shows that in the mixed group one subgroup of learners expressed higher variance of the fluctuations of the MMS amplitudes at the start of the experiment. After the second block of trials, the levels of MMS-amplitude variance systematically decreased, eventually converging to the much lower level of the other subgroup. As such, this subgroup expressed a higher bandwidth of variance values than the other subgroup. The subgroup with the lower bandwidth of variability, started out with much lower variance and remained in that regime throughout the 8 blocks of the experiment. This was the case for both target types. Furthermore, this range of variance was comparable to those observed in the purely random or purely correlated cases (shown in Figure 3C-F.)

**Figure 3.**
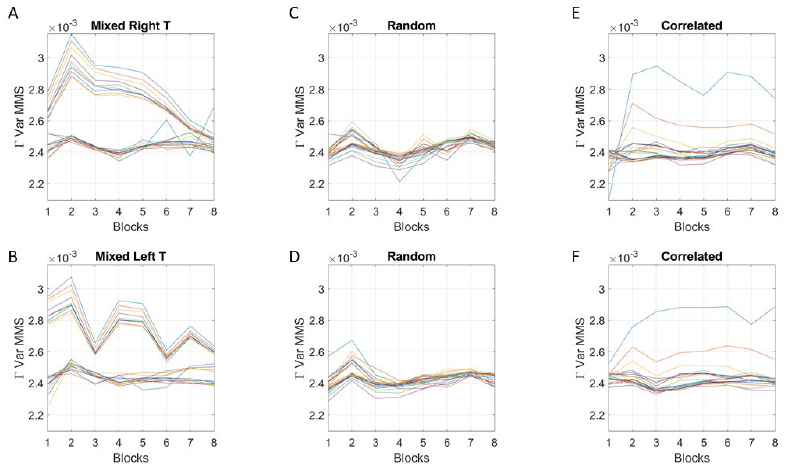
Analyses of the empirically estimated Gamma variance parameter block by block. (A-B) two subgroups in the mixed group are revealed for right and left oriented targets (each curve represents the trajectory of a participant within the group.) The subgroup with lower variance and narrower bandwidth of values throughout the experimental session separate from those in the subgroup with high variance and broader bandwidth of values. However, both subgroups converge to similar variance levels towards the 8th block of learning. Target types show different trajectories but similar convergence trend. (C-D) random group shows similar levels of variance and stable learning throughout the experimental session, as does the correlated group (E-F) (with two outliers.)

The two subgroups of the mixed condition did not differ in reaction times or accuracy, suggesting that all participants were able to reach the same online performance. Instead, they were differentiated by their explicit knowledge of the regularity imbedded in the task (Figure 4). Note that participants were not informed of the regularity. To probe whether they had nevertheless gained explicit knowledge of the repeating layouts, following the search task, participants completed an explicit memory test. A memory score was computed for each participant (see methods for details). The subgroup with broader bandwidth of variability showed low test scores (10 subjects, *M*=0.94, *SD*=0.4), thus exhibiting less explicit knowledge of the regularity. In contrast, the subgroup with the narrow, steady bandwidth of variability, gained a higher level of explicit knowledge, as reflected in higher explicit memory test scores (13 subjects, *M*=1.52, *SD*=0.75) (*p<0*.*01* nonparametric Wilcoxon ranksum test). Figure 4.

**Figure 4.**
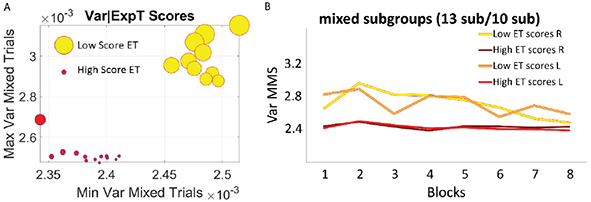
Self-emerging subgroups in the mixed group are differentiated by the scores of the explicit memory test, which probed their explicit recollection of the regularity that they experienced during the session. (A) The horizontal axis comprises the minimum value of the variance, while the vertical axis comprises the maximum value of the variance for each participant. Thus, the graph depicts the full range of variance values. The size of the marker is proportional to the explicit familiarity test score and the color represents the subgroup, with no overlapping between the two sets of participants. (B) Empirically estimated Gamma variance parameter unfolded block by block as in Figure 3AB, for the two subgroups of the mixed condition. The group with less explicit knowledge (lower scores on the explicit memory test) starts out with higher variance of the fluctuations of the MMS amplitudes, eventually converging to the much lower level of the subgroup that showed higher explicit knowledge of the regularity.

For completeness, the memory scores of the correlated group were also examined. Overall, memory scores (*M*=1.37, *SD*=0.9) were like the scores observed in the higher memory subgroup in the mixed group. This is consistent with the similar stochastic learning signatures of the correlated group and the high memory subgroup (Figure 3). As the regularity in the correlated group was highly reliable, with layouts repeating on all trials, it seems that all participants reached some minimal level of explicit knowledge, therefore no subgroups emerged.

**Global analyses** of the stochastic signatures derived from pooling trials across all participants allowed us to examine the distributions of the shape parameter describing the empirically estimated parameters of the continuous Gamma family of distributions. We reasoned that the distributions of the shape parameter, dynamically shifting as the system learned, could help us understand the differences in the mixed group with respect to the random and correlated groups. This is so because the continuous Gamma family spans distributions of different shapes. In prior research, we have empirically characterized these ranges of PDF shapes. They reflect different degrees of randomness and different levels of noise. We refer the reader to methods Figure 10D whereby we explain the empirical meaning of the Gamma parameter plane. We have empirically characterized these in human biorhythms registered across different developmental stages and different disorders of the nervous systems.

At the leftmost extreme, when the Gamma shape is 1, we have the special case of the exponential distribution. This is the case of a memoryless random process whereby events in the past do not inform more about events in the future than current events would. The information is coming from the *here and now*. Such distributions are ubiquitous in the motor code at the start of neurodevelopment ^16,29,30^. They however transition to distributions with heavy tail around 4-5 years of age, when the system is (on average, universally) mature enough to start schooling, receive instructions, and sustain longer attention spans ^16,17^. By college, age these distributions are Gaussian like. These shapes are at the other extreme of the shape axis on the Gamma parameter plane (Methods Figure 10D.) Prior work has revealed that in systems where maturation is compromised (*e*.*g*., autism across the lifespan) these global signatures remain in the exponential range, randomly relying on the here and now and manifesting excessive noise ^18,31^. Excess noise and randomness in the motor code is expressed in voluntary ^16-18,22^, involuntary ^32-37^ and autonomic ^38,39^ modes of motor control. These features prevent the acquisition of a predictive code in neurodevelopment, or it weakens it in system with neurodegeneration ^23,32,40-42^.

Figure 5 reveals this pattern for the evolution of the Gamma shape parameter. Among the moments of the distributions of the shape parameter, the variance reveals the separation of the mixed group from the random and from the correlated groups (Figure 5A). Furthermore, a distinction is also observed for the mean parameter of the distribution of Gamma shapes (Figure 5B). As such, the signal to noise ratio, SNR *i*.*e*., the mean divided by the variance (which in the Gamma family coincides with 1/scale parameter) shows the highest signal content for the mixed group (Figure 5C). For both the correlated and random groups, the distribution shape has an increasing trend, consistent in both cases for the right and left oriented targets. However, in the mixed group, there is an initial increase in the shape that decreases and stabilizes by the 4^th^ to 5^th^ block, at much lower values of the variance, so that the SNR of the mixed group is much higher than that of the random or correlated groups. This elevated SNR indicates that the mixed environment is much more effective for learning than environments that contain purely random or purely correlated trials alone.

**Figure 5.**
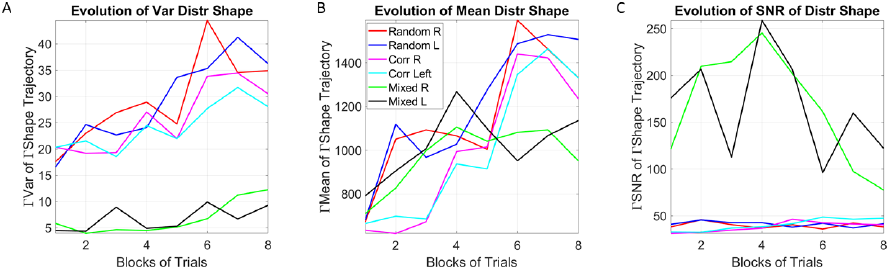
Global learning evolution shows the unstable environment (mixed group) to provide the most efficient conditions for learning, as indicated by the highest SNR. (A) Tracking, block by block, the empirically estimated variance of the distribution of gamma shape values obtained from the fluctuations in MMS amplitudes for each type of stimulus and target. Correlated and random groups trend upward with a steeper rate for correlated, while the mixed group stabilizes after ½ the session. The variance separates the correlated and random groups from the mixed group, with a marked reduction on the variability of distribution shapes and an overall trend to increase the variability in distribution shape towards the final blocks. (B) Tracking, block by block, the empirically estimated mean value of the distribution of shape values from the fluctuations in MMS amplitudes. (C) The signal to noise ratio (mean/variance) then shows the highest signal for the mixed trials, with a downward tendency after ½ the total session.

To better understand the two types of learners that were observed in the mixed group at the local level depicted in Figure 2, we performed the same analyses as in Figure 5 but this time considering separately the trials from each subgroup that emerged from the local analyses. This analysis at the global level, considering the evolution of the distributions’ shapes, revealed that the subgroup with the lowest variance in the MMS evolution (Figure 3A-bottom panel) and the highest scores in the explicit memory test (Figure 4) bears the highest signal to noise ratio. Figure 6A shows the mean, Figure 6B the variance and Figure 6C the signal to noise ratio, as the mean divided by the variance. Higher SNR in the distribution of the shape values signifies that the change in distribution shape from lower values, tending towards the memoryless exponential regime, to higher values in the Gaussian regime of the continuous Gamma family, is conducive of a more predictive code in the error corrective state of the learning progression ^43^.

**Figure 6.**
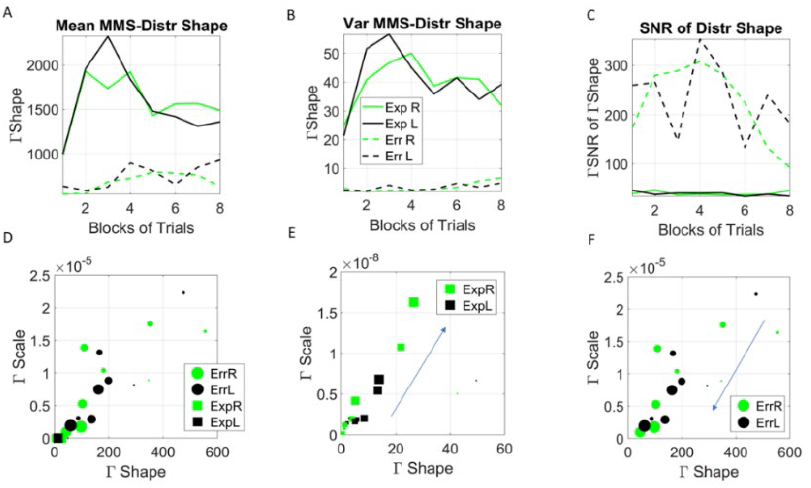
Convergent evolution across blocks of learning in exploratory *vs*. error correction modes and SNR profile across groups. (A) The evolution of the empirically estimated mean based on the distribution of Gamma shape values extracted from the MMS. (B) The evolution of the empirically estimated variance of the distribution of Gamma shape parameters shows similar differentiation patterns between subgroups of error corrective and exploratory modes, with much lower rates of change than the mean. The latter gives rise to the highest SNR (mean/var ratio) for the mixed condition depicted in (C). (D) Block by block evolution of the empirically estimated shape and scale parameters of the continuous Gamma family of probability distributions. Block number is proportional to the marker size, with earlier blocks having smaller size and later blocks increasing in size. The exploratory mode is confined to the gamma shapes close to the memoryless exponential distribution, while the error corrective mode evolves from higher to lower values of the Gaussian regime of the Gamma family. Unfolding each case (exploratory E and error corrective F) shows their convergence to a regime away from the memoryless exponential and tendency to more Gaussian like distributions. This convergent global behavior is congruent with the convergent local behavior of Figure 3.

To further characterize the type of learning of the two subgroups of participants, we examined the evolution of the distribution of the shape values, as the system experienced the learning and the PDFs shifted shape. The shape parameters of the subgroup with high explicit memory scores (Figure 4AB) starts in the symmetric Gaussian range but converges towards the skewed to exponential range. This confirms the departure from a memoryless random state and the prevalence of a state where higher predictability is attained under higher signal to noise ratio in panel 6C. This is consistent with the profile expected in error correction learning. In the error corrective mode, a goal, that is, the minimal knowledge that some regularity is present within the incoming stream of information, has been detected, so that the system knows what it needs to learn (*i*.*e*., has self-discovered the goal).

For the subgroup with low explicit memory scores (Figure 4AB) the same analyses showed a markedly different learning profile: the values of the shape parameters for these participants varied confined within the exponential range of the continuous Gamma family, *i*.*e*., with shape parameter value close to 1, for the exploratory mode. This is shown in Figure 6D (squares), where the contrast with the subgroup with high explicit memory score (*i*.*e*., error corrective cohort) (circles) is noted (Figure 6E zooms into the exploratory scatter). This profile corresponds to an exploratory phase of the learning, occurring within a time when the system is trying to evaluate the presence of predictive information. Such wondering behavior gives rise to more variability (as depicted in Figure 3A, 4B). In this case, during the initial trials, the system is sampling information in the *here and now*, not yet relying on prior information, but rather examining the information as it comes, trial by trial. Presumably, as minimal explicit knowledge is obtained, this profile gradually converges into a learning profile consistent with the discovery of some regularity. Then, the signature of an error correction account of learning emerges. This inference is suggested by the unfolding trajectories of Figure 3A-B for the local level, but also further refined for the global level of Figure 5.

As shown by Figure 3 at the local level, here at the global level, the two subgroups (exploratory and error-corrective) converge to comparable levels of variance (of the MMS amplitude in Figure 3A-B and of the distribution shape in Figure 6E-F), where the shape of the marker is proportional to the block number and the arrow marks the direction of the flow (from lower to higher shape values in Figure 6E, and from higher to lower shape values in Figure 6F.)

At a global timescale (*i*.*e*., entire experimental session) we assessed the change in stochastic variations of the signals over time. To do so, we examined the evolution of the fluctuations in the change of distributions using the Earth Movers Distance (EMD) metric (see trajectories in the supplementary Figures 1-3). We compared from trial to trial, and block to block, across participants, the fluctuations in the amplitude of the change in distributions (as measured by the EMD.) We also assessed the rate of the change in peaks (inter peak intervals related to the timing of the overall global process.) These parameters amount to a “speed temporal profile” of the PDFs as they shift stochastic signatures per unit time on the Gamma parameter plane. The analyses revealed that the system clearly distinguishes the rates at which the distributions change shape from the random to the correlated groups and between those and the mixed group. Figure 7A shows this on the log-log Gamma parameter plane where each point with 95% confidence intervals, represents the performance for the right target (left not shown for simplicity but has similar patterns.) The corresponding PDFs for both right and left oriented targets are shown in Figure 7B. We can appreciate that the mixed case yields the most predictive shifts in distribution change, with the highest shape value and the highest signal to noise ratio (*i*.*e*., the lowest Gamma scale value.) Furthermore, these rates of change in the two subgroups of the mixed case, clearly distinguish the left from the right oriented targets, with comparable rates of shifts in distribution shape for the exploratory and the error corrective subtypes. These are shown in Figure 7C (estimated Gamma parameters) and 7D (corresponding Gamma PDFs.) These comparable shifts in distributions hint at a smooth process whether the system is curiously wondering in exploratory mode, or aiming for a targeted goal, in error corrective mode.

**Figure 7.**
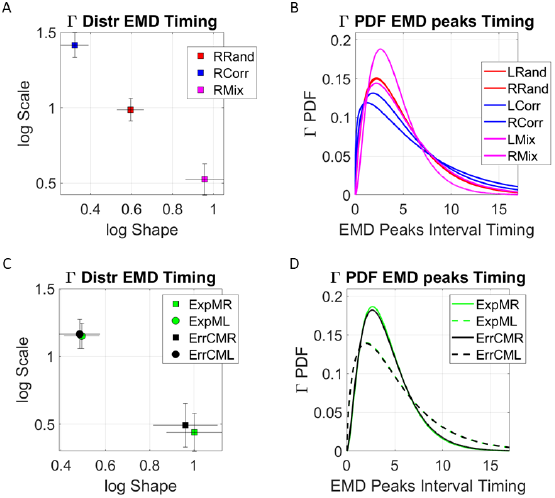
Global dynamics. Unfolding the global rate of change in distribution shapes, as the system transitions from PDF to PDF, using the EMD to ascertain distribution differences from moment to moment. (A) Right target case is shown for the three groups with 95% confidence intervals for the empirically estimated Gamma shape and scale parameters. Each point represents a different distribution. Here the mixed group shows the maximal values of log shape (Gaussian) and SNR (1/log scale). (B) The PDFs corresponding to the maximum likelihood estimation (MLE) distributions in (A). (C) Investigating the differentiation between targets for the two subgroups of the mixed condition at the global level reveals similar rate of change in the interpeak intervals, suggesting smooth transitions in both exploratory and error corrective cases, with clear target differentiation. (D) Corresponding PDFs for (C).

Finally, we examined, block by block, the evolution of the hubs in the EEG leads as nodes of the network. This dynamical shift in activity is shown in Figure 5 of the supplementary material. In general, we find the lead P1 (located over the left parietal area) as the most heavily recruited node in both target orientations and for both learning modes. However, in the exploratory mode, with higher variance, there is a more distributed recruitment of hub nodes across the recorded area (posterior part of the head) with marked differentiation for left *vs*. right oriented targets. We caution that our analyses were constrained to a subset of leads (see methods) owing to avoidance of artifacts from eye and jaw motions, along with other motivational prior work ^44^, and refrain from stating anything about sources underlying the leads’ activities. Instead, we report the changes in leads representing hubs that self-emerge from the activities across the central and posterior leads that we included in these analyses (for details see Figure 5 in the supplementary material). Beyond extracting dynamic information from the hub channel for each participant, we did not examine other patterns of brain activities pertinent to each learning mode, though we know that the posterior parietal cortex is itself a hub linked to predictive codes that emerge during the learning of novel situations ^45-47^. This type of research is also warranted, to learn more about causal network interactions in the space of leads.

## Discussion

This study evaluated online dynamics of statistical learning using a new approach that relies on gross data as the main measure of interest. Specifically, we characterize the evolution of the learning process using the moment-by-moment standardized fluctuations in the peak amplitude and timing of an EEG signal recorded while participants performed in a visual search task. Depending on the group, and unbeknown to participants, the task contained a reliable regularity (correlated group), no regularity (random group) or a relatively unreliable regularity (mixed group). We found that statistical learning is a highly dynamic and stochastic process, sensitive to the reliability of the incoming information. Moreover, we discovered that embedded in the gross data, traditionally discarded as superfluous noise under assumptions of normality, lies a code that describes different modes of learning. Based on our stochastic characterization of the learning phenomena at different local *vs*. global time scales, we equate this distinction with two fundamentally different types of processes. These are the commonly studied error correction type linked to stimulus regularity and the newly characterized, exploratory process likely reflecting self-discovered surprising contextual variations. To aid interpreting these results, we leverage prior research on the broad characterization of human biorhythmic activity and reframe this problem from the vantage point of neuromotor control ^16,21,29,33,36,37^.

Two main results emerged from the current analyses. First, we show that unstable environmental conditions (*i*.*e*., mixed group) provide the most opportunity for learning, as characterized by higher signal to noise ratio on both the global and local levels of analyses. Next, we show that on an individual basis, this unstable environment may give rise to different learning profiles: within the mixed group, two subgroups of participants self-emerged from the analyses. For one subgroup, the learning profile corresponded to error correction mode from start to finish. However, for the second subgroup, the learning profile reflected an early stage of exploratory learning which later converged into the error correction mode. Crucially, despite no *a priory* assumption, these two classes of learners were differentiated behaviorally by their level of explicit knowledge of the regularity in the task. We now turn to discussing each of these results in detail, while considering their implications on our understanding of statistical learning in general.

### Unpredictable environments provide more opportunity for learning, corresponding to a more efficient learning process than predictable environments

When comparing the stochastic signature of learning within an unstable environment (mixed group) with stable conditions (correlated and random groups), the process proved to be less stationary, more predictable in nature, and was characterized by higher signal to noise ratio. These characteristics suggest that complex environments provide higher opportunity to learn than reliable environments. Moreover, within our theoretical framework, higher signal to noise ratio corresponds to more efficient learning. These results are consistent with neuroimaging studies that have identified brain systems that track uncertainty in a curvilinear U-shaped function ^3,48^. Within these systems, full randomness or full regularity are alike in terms of informativeness and provide less information than the mixed case. As such, these systems seem to be especially sensitive to tracking relatively unreliable information in the environment.

Given that the real world is indeed complex, with our cognitive system continuously bombarded with variable regularities, it seems natural that we should be more attuned to learning under relatively unreliable conditions. However, suggesting that learning under such conditions is more efficient may seem to contradict the behavioral pattern observed in the data: participants in the mixed group reached slower RTs than both in the random and correlated groups (for a detailed account see ^15^. To resolve this issue, one must bear in mind that efficiency of learning is not necessarily manifested in online performance. That is, more complex learning conditions may hinder online reactions, but be beneficial for the long term. We propose that to gain further insight on statistical learning, future studies should combine the methods introduced in the current work with experimental designs that involve changing regularities online and considering multiple sessions of learning. Indeed, such designs are becoming common within the field ^49-51^. However, so far, they lack the perspective of evaluating the online evolution of the learning process, which is enabled by the methods used in the current work.

### Exploratory vs. Error correction modes differentiated by explicit knowledge of the embedded regularity

For a subgroup of participants, the unstable environment (mixed group) triggered an initial stage of exploratory learning. During this stage, the stochastic signature of the process reflected a memoryless type of learning, suggesting that the system was not relying on prior knowledge but instead gathered as much information as possible from the “*here and now*”. Presumably, this exploratory stage was elicited by the high levels of surprise in an environment that contained rules that were not followed consistently over time. Crucially, this subgroup also exhibited low scores in the explicit knowledge test. For participants that showed higher level of explicit knowledge, the stochastic signature reflected an error correction mode of learning from the beginning of the task. Given this behavioral differentiation, it seems that the transition from exploratory behavior into error correction depends on some minimal level of explicit knowledge that needs to be obtained. This conclusion contradicts the current assumption that both explicit and implicit statistical learning always reflects error correction ^3,4^. For instance, within theories arguing that both explicit and implicit learning systems operate simultaneously (*i*.*e*., dual-system approach), it has been suggested that during a learning episode, implicit associative learning occurs initially, which leads to the formulation of predictive “wagers” that steadily become more correct, leading to explicit awareness of the learned patterns ^52^. The initial stage of exploratory, memoryless sampling from the perceptual input that has emerged from our analyses has so far been overlooked.

The new methodology introduced in this work is grounded on deliberate *vs*. spontaneous movement classes ^19^, with different classes of temporal dynamics. Framed in this way, the error correction code would correspond to the deliberate movements intended to a goal. Such movements are well characterized by paths that can be traversed with different temporal dynamics and remain impervious to changes in speed ^21,53-57^. Within such learning, the path to the goal is independent of how long it takes to attain it and remains stable despite the moment-by-moment temporal structure of the stimuli, which must be learned and transformed into physical, motoric action ^53,55,58^. This invariance is akin to timescale invariance in models of temporal learning, strongly supported by empirical data ^59,60^. In contrast, exploratory learning, would correspond to the class of spontaneous movements, *i*.*e*.., highly sensitive to contextually driven variations in temporal dynamics of the stimuli ^19,21,24^. These different dynamics can be distinguished in the variance profile of the learners in the mixed group of Figure 3A-B. They respond dynamically different across blocks, depending on target type. In this sense, exploratory trajectories with higher variance, lower explicit memory scores and fundamentally differentiating target responses, are contextually more informative than error correcting trajectories. During this exploratory mode, *all events are equally probable*. The system samples without restriction. This mode may increase the chances of surprise, grabbing the system’s attention to some context relevant events, perhaps self-discovering the transition toward regular, ever more systematic states that may eventually result in a desirable, stable goal to then guide error correction mode. Such smooth transition between modes was evidenced in the convergence of the variance profile to a common regime.

Information about context may come in the memoryless exponential regime that prevails in the early stages of learning for the subgroup in the mixed condition. At the global timescale, this early learning with distributions of exponential shape, was accompanied by high SNR in those early trials. This feature differs from the type of memoryless exponential signature found in autism ^16^. In the case of autism, the presence of the memoryless exponential distribution, especially during instructed learning, is paired with low SNR ^16-19,21^. This high random noise regime in autism adds a type of uncertainty that interferes with the overall learning process. It leads to high levels of stress and anxiety during prompt-mediated learning ^22,33,36,37^. In this sense, we posit that the type of learning observed here, *i*.*e*., characterized by low NSR in the memoryless exponential phase, may be that of a curious, wondering system, welcoming surprising events that lead to more spontaneous exploration (*e*.*g*. present as well in the spontaneous hand motions of pre-verbal autistics who implicitly, without instructions and largely beneath awareness, self-discovered the goal of a task ^20^ and especially, present much earlier in neonates that learn how to control their motions during the first three months of life ^30^.)

We propose then, to trace back the newly characterized exploratory mode to the neonatal stages of learning. Such stages appear prior to the maturation of perceptual systems and are guided by endogenous bodily fluctuations that the infant senses from self-generated movements (likely heavily involving central pattern generators). To that end, we cite how neonates learn, perhaps supporting our idea that humans’ mental strategies and the different learning modes discovered here, are embodied, grounded on the types of learning that we phylogenetically transitioned through during early infancy ^25^. Studies of infants exploring an environment where the mother serves as an anchoring reference place, find that the babies explore using interleaving segments of progressive movements with lingering episodes ^6^. They confirm that such exploratory behavior is homologous across species and situations ^5,6^. Furthermore, a recent study from the statistical learning domain demonstrated that infants prefer to attend to events that are neither highly unpredictable nor highly predictable ^61^. The authors suggest that this effect is a characteristic of immature members of any species, that must be highly selective in sampling information from their environment to learn efficiently. We add to these interpretations and suggest that infants attend to relatively unpredictable environments because these are ideal for the exploratory behavior that dominates early stages of surprise- and curiosity-driven motor learning in neonates ^30^ and infants ^16^.

Across early stages of life, when altricial mammals generally mature their somatic-sensory-motor systems ^62^, human infants acquire a stable motor percept. As they undergo motor milestones (myelination, acquisition of motor, and sensory maps, *etc*.), the families of PDFs that are empirically estimated from purposeful movements, transition from memoryless exponential to highly predictive Gaussian ^16,20,30^. However, given our results, it appears that the exploratory type of learning is preserved throughout adulthood, and that there are conditions in which this exploratory, memoryless learning emerges on demand, and is likely extremely advantageous. An open question is, when is this type of learning beneficial? One possibility is that it supports flexibility within the system, as it provides it with a range of information that would have been missed had an exploratory mode not been evoked. That is, in changing, unstable environments, it may be best to initially gather as much information as possible, before committing to an error correction, goal-targeted mode. This direction, which is beyond the scope of the present work, may be tested by examining whether exploratory periods emerge during processes that require flexibly extending an existing solution to new context, known in motor control as transfer and generalization ^63-65^. Generalization and transfer have been previously studied in exploratory mode ^20^ but such studies are rare. This research may bear important implications for clinical programs that are currently grounded in animal models of conditional reinforcement that do not address the possible benefits of an exploratory mode of learning.

Note that the presently uncovered learning modes are different from known computational models of reinforcement learning whereby exploration *vs*. exploitation modes are used to model learning through rewards ^66,67^. In these models, exploration and exploitation learning are differentiated by the type of reward, which is either intrinsically obtained, or extrinsically provided. Yet for both modes, the learning is best described by error correction, as both in the exploitation and exploration modes the system considers previously gathered information and moves towards a desirable configuration. The exploratory mode proposed in this work is a memoryless process that does not pursue a goal. In this mode, the system is sampling information as it comes, casting the widest net possible, before enough predictive information is gathered at random, thus prompting the transition into the error correction mode. Related, are recent models of human and machine learning that emphasize the role of curiosity within the learning system ^68,69^. These models suggest that the causal environment determines when curiosity is driven by novelty or by prediction errors. In an environment where the past and future occurrences of stimuli are independent of each other, the optimal solution for gaining a future reward is to explore novel stimuli. This novelty mode, that has been referred to as novelty-error-based ^68^, and the standard prediction-error-based approaches have at their heart the same computational problem: optimize with respect to an error that depends on a given target goal, while using prior information. The exploratory mode suggested in this work is computationally different from the error correction mode, as it does not operate with a goal in mind and gathers information initially on a memoryless way, regardless of a goal or reward.

We argue that to characterize learning properly, this additional type of endogenous, curios exploration should be incorporated into future models. Furthermore, we posit that this exploratory mode described here, scaffolds the emergence of *spontaneous autonomy*, different from deliberate autonomy (*i*.*e*., derived from targeted error-correction.) It will be critical to include exploratory learning in the future design of autonomous robots/agents. This type of autonomy and its accompanying sense of agency can be realized through the autonomous self-discovery of the relationships between actions and their consequences. The latter leads to the sense of action ownership and to the volitional control of physical acts that are congruent with one’s own mental intent. Only then, after acquiring this congruency, can others understand one’s intent.

We have in summary shown the dynamic nature of statistical learning, the rich stochastic signal embedded in fluctuations that are traditionally treated as gross data and the differential nature of different learning modes. Investigation is warranted on whether these results generalize to other statistical learning paradigms, and to the acquisition of predictive information in learning in general. Of particular interest, are questions of individual differences, and the degree to which the exploratory and error correction learning modes may be differently recruited on demand by the same learner under different contexts. We here offer methods that allow to investigate these and many new questions in future statistical learning research from the perspective of the nascent, developing motor systems and their richly layered dynamic and stochastic motor percepts.

## Methods

### Participants

Data from 70 participants (48 female, mean age, 23.7) was analyzed in this study: 24 in the random group, 23 in the correlated and mixed groups. There were no differences in age or gender between the three experimental groups. Two participants (one in the mixed group and one in the correlated group) from the original study were excluded due to a bug: their EEG recording started late, missing the first few trials. This issue was not significant in the original analyses of the data as it relied on epochs of the signals and their averages ^15^. As we now focus on continuous data analyses, we decided not to include these two subjects.

### Stimuli and Procedure

All participants gave informed consent following the procedures of a protocol approved by the Ethics Committee at the Tel Aviv University. Next, participants received a general explanation regarding EEG experiments and completed three tasks: visual working memory (VWM) capacity assessment, visual search task and a surprise explicit memory test. EEG was recorded only during the visual search task. In the present work we focus on the visual search task and the surprise memory test. We did not find any connection between the VWM task and the present results. As such we do not describe this task further. A detailed account can be found in ^15^.

Stimuli in the visual search task and the familiarity test were white T’s and L’s (see Figure 1B in the main text). All stimuli were made up of two lines of equal length (forming either an L or a T). From a viewing distance of approximately 60 cm, each item in the display subtended 1.5° × 1.5° of visual angle. All items appeared within an imaginary rectangle (20° × 15°) on a grey background with a white fixation cross in the middle of the screen (0.4° × 0.4°). Targets appeared with equal probability on the right or left side of the screen.

### Visual search task

Participants searched for a rotated T (target) among heterogeneously rotated L’s (distractors) while keeping their eyes on the fixation cross. Each trial began with the presentation of a fixation cross for 2100, 2200, or 2300ms (randomly jittered) followed by an array of one of two possible targets (left or right rotated T) among seven distractors. Participants were instructed to press a response key corresponding to the appropriate target as fast as possible. Depending on the group, the visual search contained the consistent mapping condition, the random mapping condition, or both. For the consistent mapping condition, spatial configurations of targets and distractors were randomly generated for each participant (8 layouts for the mixed group and 16 layouts for the correlated group). In the random mapping condition targets and distractors appeared in random locations throughout the task with the exception that for the mixed design group, targets in the random condition could not appear in the same locations as targets in the consistent condition. The order of layouts was randomized every 16 trials (in the case of the mixed group 16 trials correspond to 8 consistent and 8 random trials presented in a random order). In all conditions the identity of the target (left or right rotation) was chosen randomly on each trial and did not correlate with the spatial regularity. Participants completed 512 trials in the experiment.

### Explicit memory test

Participants were not informed of the regularity in the task. Upon completing the visual search task, participants in the mixed and correlated groups (groups that performed in a task that contained regularity) completed an explicit memory test, designed to reveal explicit knowledge of the regularity: participants saw the layouts that were presented to them during the search task mixed with new, randomly generated layouts. For each layout participants had to indicate whether they have seen the layout during the visual search task or not. We then computed a memory score (hit rate/false alarm rate) that is considered to reflect each participant’s explicit knowledge of the regularity. Higher scores correspond to better explicit knowledge. For more details, see ^15^.

### EEG recording

EEG was recorded inside a shielded Faraday cage, with a Biosemi Active Two system (Biosemi B.V., The Netherlands), from 32 scalp electrodes at a subset of locations from the extended 10–20 system. The single-ended voltage was recorded between each electrode site and a common mode sense electrode (CMS/DRL). Data was digitized at 256 Hz (For a more detailed account see ^15^. As we rely on continuous recordings, without taking averages of data epochs, we focus on the electrodes that do not reflect strong eye muscle activity either through blinking or the jaw movement. The analyzed subset Fp1, Fp2, AF3, AF4, F3, F4, F7, F8, Fz, FCz, C3, C4, Cz, T7, T8, P1, P2, P3, P4, P5, P6, P7, P8, Pz, PO3, PO4, PO7, PO8, POz, O1, O2, and Oz), includes all of the electrodes that were previously analyzed (P7, P8, PO3, PO4, PO7, PO8, C3, C4) ^14^. Only correct trials were included in the analysis.

Data was analyzed on two levels: unfolded for 8 blocks and overall. In both cases the data was not averaged, but as described later in this section evaluated continuously. Each block corresponded to approximately 4 minutes of recording. The blocks were defined by the breaks in the task. During a break, participants could move freely, drink, and rest their eyes. If needed, EEG recording issues were addressed during the breaks.

### EEG preprocessing

We use the EEGLAB PREP pipeline 70 to clean the EEG signals. Furthermore, we do not use the signals from the break times within the session, when the participants were moving and disengaged from the task. We use the precise landmarks of the study to separate the correct *vs*. incorrect trials, the trials within each block and focus on the experimental epochs of interest as they continuously evolved within the session, gathered by target orientation type (left *vs*. right) and by group (mixed, random, correlated.) These analyses are centered on the responses from the correct trials.

### Micro-movement Spikes

Figure 8 shows the transition from a continuous (analogue) signal to discrete spikes (digital) signal. We coin this signal the micro-movements spikes MMS. This is a general datatype that standardizes the original timeseries waveforms from each participant. We do so by scaling out anatomical differences, while preserving the original ranges of the data ^71^.

**Figure 8.**
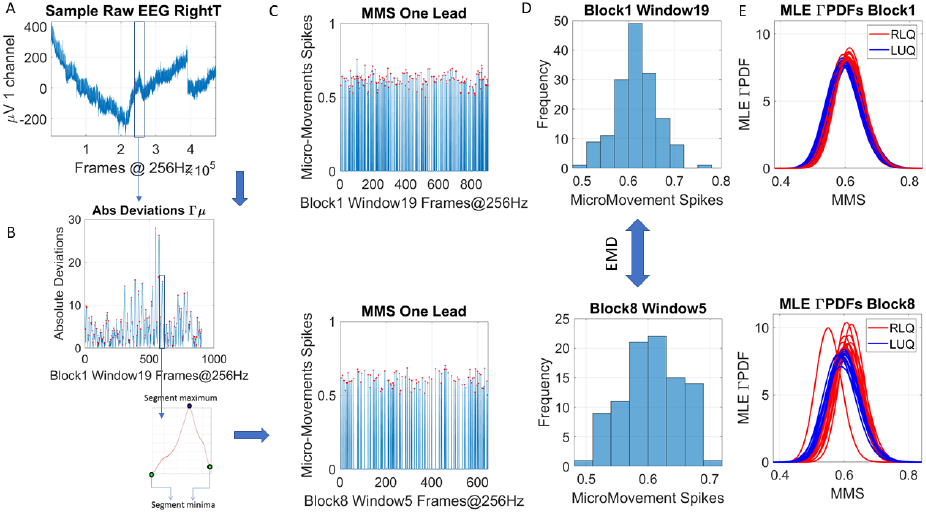
Transforming continuous analogue signals to digital spikes: micro-movement spikes MMS. (A) Sample electroencephalographic signal from one channel, zooming into one segment. Sweeping through the signal, windows of 5 seconds with 50% overlap are taken to scale each peak value deviated from the empirically estimated mean (μV). (B) To that end, for each participant, the original peaks are used to empirically estimate the mean amplitude across the session, and obtain, for each point in the time series, the absolute deviation from the empirically estimated mean. This series of fluctuations are then used to scale out possible allometric effects from *e*.*g*., anatomical head differences, using equation 1 in the methods. As in the inset, each peak in the segment (segment maximum) is surrounded by points between segment minima. Equation 1 is used to obtain the unitless, standardized MMS. (C) The resulting unitless quantity is plotted as a series of MMS for two sample states in some window of blocks 1 and 8. (D) The peaks (red dots) are gathered into a frequency histogram to obtain the difference, from window to window (block by block), using the earth movers’ distance, a similarity metric used in transport problems. We then obtain the amount of effort that it takes to transform one frequency histogram into the other. (E) Using maximum likelihood estimation (MLE) the best continuous family of probability distributions fitting each histogram is obtained. In this case, the shape and scale parameters of the continuous Gamma family are estimated for different states of the stochastic process (explained below), as they transition from random noise to predictive signal, increasing the signal to noise ratio of the MMS.

The notion of MMS has been patented by the US Patent office ^23^. To obtain it from any biorhythmic data, we take the peaks and valleys of the original waveform and perform a transformation that nevertheless preserves the original ranges of the data. In this case, the peaks, and valleys of the continuous EEG waveform (μV) are obtained (*e*.*g*., the waveform from a lead is shown in Figure 8A.) Then, the values of the peaks are gathered into a frequency histogram. To that end, we explored windows between 1 and 5 second-long (with 50% overlap) and settled on 5 seconds as the minimal time unit that gave us acceptable 95% confidence intervals in the empirical estimation process requiring close to 100 peaks or more. Note that other sizes of window above 5 seconds are also possible, but difficult to interpret physiologically as we move away from sub-second time resolution. In general, we caution that the sampling resolution of the sensors in use will determine the size of the window of choice for empirical estimation of the distributions (and their shifts), as it will determine the number of peaks read out from the biorhythmic activity per unit of time. One should consider that the unit of time is thus empirically determined for at least ∼100 peaks/unit time. In the Figure 8B example 97 peaks (red dots) extracted from the window (from 1,280 points, 256 points/second x 5 seconds, whereby 97 depart from the estimated Gamma mean, explained next.)

Using maximum likelihood estimation (MLE), we approximate the mean (in this case the Gamma mean, as the continuous Gamma family of probability distributions best fits the histograms in an MLE sense, compared to the log normal, the normal and the exponential distributions.) We obtain the absolute deviation of each point in the original EEG time series, from the empirically estimated mean. These values are then used to build an orderly series of values and its peaks and valleys used to scale out allometric differences owing *e*.*g*., to anatomical differences in head circumference. The local peak (maxima) of these series of fluctuations is then divided by the sum of its value and the averaged values of points between the two local minima surrounding it (see equation 1 below and Figure 8B.)

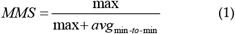

This information is then plotted as in Figure 8C reflecting the unitless, standardized MMS describing the minute fluctuations in the original waveform away from the empirically estimated mean. We group the peaks (marked by the red dots) into a frequency histogram and use the Earth Mover’s Distance, EMD ^72,73^, to measure the difference between the two distributions.

### Cross-Coherence Analyses and Network representation

The cross-coherence between two times series (assumed to be the realizations of unknown stochastic processes) is defined as the cross-spectral density between the two series normalized by the product of their auto-spectral densities ^74^. We use cross-coherence to quantify the similarity between the MMS series of any two leads in the frequency domain (*e*.*g*., two leads in Figure 9A and their original waveforms in Figure 9B are used to explain the analytical pipeline.) The frequency histograms are shown in Figure 8D for two different blocks and different windows. Each window comprises the MMS derived from 5 seconds worth of data sampled at 256Hz. The PDFs thus estimated are shown for the windows and Blocks 1 and 8 in Figure 8D. *Mean frequency* of a spectrogram *P*(*f*) is calculated as:

**Figure 9.**
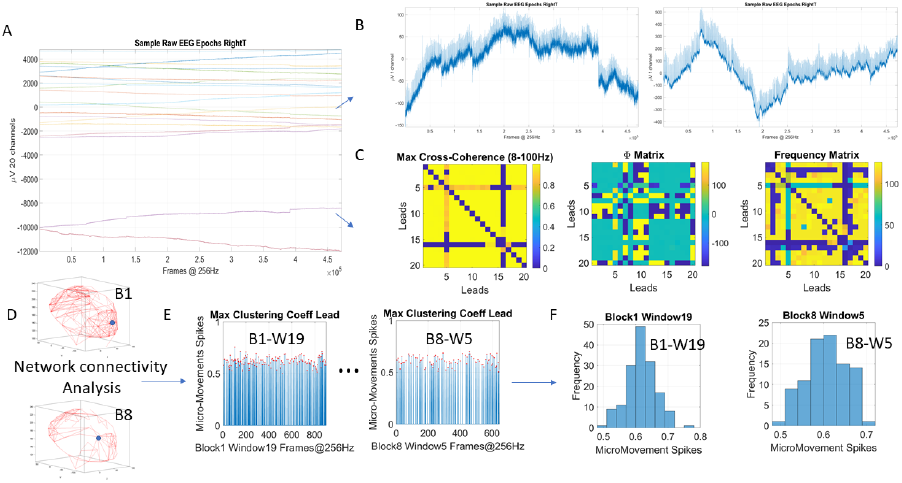
Pipeline of network connectivity analyses to select hubs for stochastic analyses. (A) The electroencephalographic (EEG) activities from twenty channels and approximately ½ hour is sampled at 256Hz. (B) Two sample leads are used to instantiate the analyses. The pairwise cross-coherence is obtained. (C) For each pair, the maximal cross-coherence is obtained, with corresponding phase and frequency values at which the maximum is attained. These build three 20×20 matrices to parameterize (for each window and across each block of the session) the activity and build adjacency matrices (using the maximal cross-coherence matrix.) (D) The maximum clustering coefficient in each window is obtained, here represented in schematic form for Blocks 1 and 8 (using windows 19 and 5 for visualization purposes.) (E) The MMS are obtained, and the frequency histograms (as in Figure 8) used to obtain, pairwise, the EMD matrix.

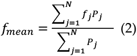

Where *f*_*j*_ is the central frequency of the j-th bin of the spectrum and *P*_*j*_ the corresponding value of the power spectral density. N is the total number of bins ^75^.

Upon pairwise comparison, we then identify the frequency for which the cross-coherence function is maximized. Figure 9C shows the three 20×20 matrices that serve as a parameterization of the signals. These include the maximal cross-coherence matrix, the phase matrix with each entry representing the value of the phase (or of the frequency, respectively) at the maximum cross-coherence value.

The maximal cross-coherence matrix is used as an adjacency matrix to build a weighted undirected graph representation of a network. Network connectivity analyses then are used to obtain the maximum clustering coefficient representing the hub within each window of activity in a block. Then, the block-by-block activity is tracked in the hubs. For each hub (represented in Figure 9D shifting from Block 1 to 8) we obtain the MMS (Figure 9E) and follow with the computation of the frequency histogram and EMD metric (as explained in Figure 8). These are used to empirically estimate the stochastic process described below ^76^.

### Empirical Statistical Estimation

The normalized peaks from the MMS are used to plot a frequency histogram (*e*.*g*., Figure 9F) for each window within a block and across blocks in each session. We then fit a PDF using MLE (*e*.*g*., Figure 8D). This is done through the estimation of the Gamma (*a*) shape and (*b*) scale parameters of the continuous Gamma family of probability distributions. The Gamma family choice comes as a result from MLE, whereby it has been found to be the optimal means of representing MMS derived from human biorhythmic data ^29,77^. This has been the case in voluntary motions, in spontaneous motions, in involuntary motions, and in autonomic motions ^29^ derived from EEG ^46,76,78^, ECG ^38,39^, kinematics parameters ^16,33,79^ and genes expression ^44,80^.

The plane spanned by the shape and the scale parameters of each Gamma PDF derived from the MMS in each window, are then plotted with 95% confidence intervals as points along a trajectory, on the Gamma parameter plane. For example, Figure 10A shows the progression of such points (block 1 on the top panel and block 8 on the bottom panel.) Different colors represent different order, such that within a block, a distribution with high shape value (symmetric) and low scale value (low NSR) may shift to a location with low shape value (towards the memoryless exponential regime) and high scale value (towards noisy regimes.) Likewise, from block to block, these transitions occur in random order. To track the stochastic behavior of the trajectory and characterize the transitions, we quantify the amplitude of the change and its direction. First, we take the median of the shape values and the median of the scale values and draw a line across each axis, to break the Gamma parameter plane into quadrants. On a log-log scale, we then plot the points and divide them into those in the Right Lower Quadrant (RLQ) and those in the Left Upper Quadrant (LUQ) as in ^30^. This division is also tracked on the Gamma moments space of Figure 10C, where we project the points of the Gamma parameter plane as the estimated moments of the distribution and show their corresponding shifts on this parameter space. This visualization has empirical interpretability from having characterized thousands of participants across the human lifespan and across diseases and disorders of the nervous systems. This empirical interpretation is shown on panel 10D. Here we see that along the shape axis, distributions shift from the memoryless exponential to distributions with heavy tails, to symmetric distributions. Along the scale axis, we move from low NSR, to high NSR. Because there is a tight linear fit to the log-log scatter, knowing the shape, we can then infer the scale (and vice versa.) As such, we reduce the number of parameters of interest to one, as we can use one of the Gamma parameters to make inferences about the state of the process. As the stochastic signatures shift from moment to moment, we can track both the direction and the magnitude of the shift. To track the direction, we use the quadrant’s location. To track the magnitude of the shift, we use the EMD as a similarity metric (proper distance metric) that informs us of the amount of “work” that it takes to change one frequency histogram into the other ^73^, as we track window by window, the activity of the hubs within a block. This metric approach enables us to probe multiple directions of change (owing to variable stimuli) and select the direction which causes the maximal change at each step (*i*.*e*., if we were to move along a desirable target-driven gradient of an objective function as in ^53^ in the case of error correction, or just explore without a desired target.) The general formula for the PDF of the gamma distribution is shown below (equation 3), where *a* is the shape parameter and *b* is the scale parameter.

**Figure 10.**
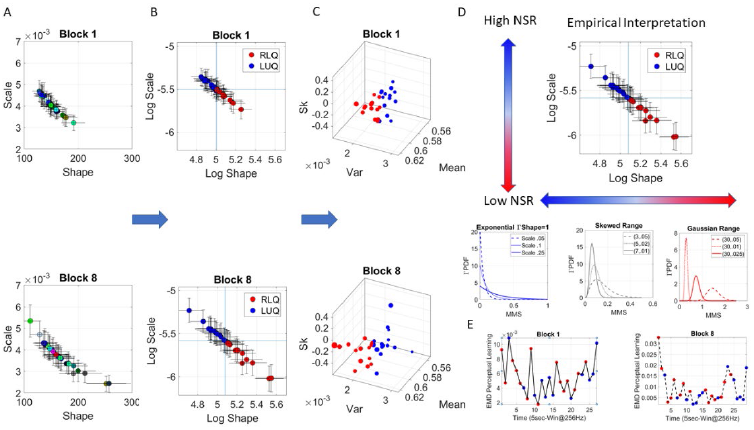
Stochastic analyses of the MMS derived from hub’s activities. (A) Upon determination of the hubs (taken window by window and block by block, across the session, the MMS of the hub lead is obtained and MLE used to determine the parameters of the best continuous family of probability distribution functions (PDFs) describing the MMS. In this case the Gamma family. The Gamma shape and scale parameters thus estimated, are then plotted with 95% confidence intervals, on the Gamma parameter plane. Window by window, and block by block, these stochastic shifts are tracked as a trajectory, whereby the magnitude of the shift and its direction are obtained. Figure shows examples of the trajectory points obtained across 5-second-long windows for blocks 1 and 8. Colors represent arbitrary order. (B) The log-log Gamma parameter plane is obtained to track points according to the quadrants spanned by the median shape and median scale, taken across each block. The Right Lower Quadrant RLQ contrasts with the Left Upper Quadrant LUQ. These have empirical interpretation. (C) The Gamma moments are obtained (see methods) to visualize the points in (B) on a parameter space whereby the Gamma mean is represented along *x-axis*, the variance along the *y-axis*, the skewness along the *z-axis* and the size of the marker is proportional to the kurtosis. The color corresponds to the direction of the shift, where the point lands, red is from the LUQ to the RLQ, or from the RLQ to itself, whereas blue is from the RLQ to the LUQ, or from the LUQ to itself. (D) Empirical interpretation of the Gamma plane and the quadrants. Along the shape axis, the distributions change from the shape *a=1* memoryless exponential to the Gaussian range, with skewed distributions with heavy tails in between. Empirically, exponential regimes are associated with immature systems, early learning in infancy and disorders or the nervous systems, if the noise to signal ratio (scale) is high. Gaussian-like regimes are associated with predictive codes, athletes or dancer’s expertise and maturity. Skewed distributions are present across maturational stages and typical learning. Along the scale axis, high values represent high noise to signal ratio (see text) which is found in neuropathy, autism, Parkinson’s disease, schizophrenia, and other disorders with specificity determined by levels of control (voluntary, involuntary, autonomic, spontaneous.) (E) The EMD is used to track the magnitude of the shift from PDF to PDF, while the direction is tracked by the quadrant landing. This curve represents the evolution of the stochastic process and serves to determine *e*.*g*., critical points of transitions for each block of the session.

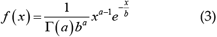

The moments (μ, σ, skewness, kurtosis) are 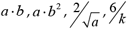 respectively. For this reason, 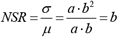 is the scale which we track as part of the evolution of the stochastic signatures. Figure 10E shows the rate of change of these transitions within block 1 (left) and block 8 (right.) The color of the dot represents the landing direction of the stochastic shift (from LUQ to RLQ, red, or from RLQ to LUQ, blue; they also represent landing on the same quadrant, red is for RLQ to RLQ and blue for LUQ to LUQ, but we could also carve out these cases separately to assess stationarity per unit time in each quadrant) while the EMD (*y-axis*) represents the magnitude of the shift. Following our empirical interpretation panel 11D, when the points are primarily red, the distributions are skewed to symmetric (towards the Gaussian range of the Gamma family.) When the points are primarily blue, the distributions are skewed to memoryless random (with *a=1* at the memoryless exponential limit.) In the former case, the signal to noise ratio (*1/b*) is higher and the process more predictive. In contrast, at the other extreme, the process is memoryless, more random and with lower SNR. It is possible to track this evolution for each participant, across trials of a block, or across blocks of a session, *i*.*e*., locally and / or globally. Figure 11 shows the global analyses used in Figure 7, further discussed in the Supplementary Figures 1-3.

**Figure 11.**
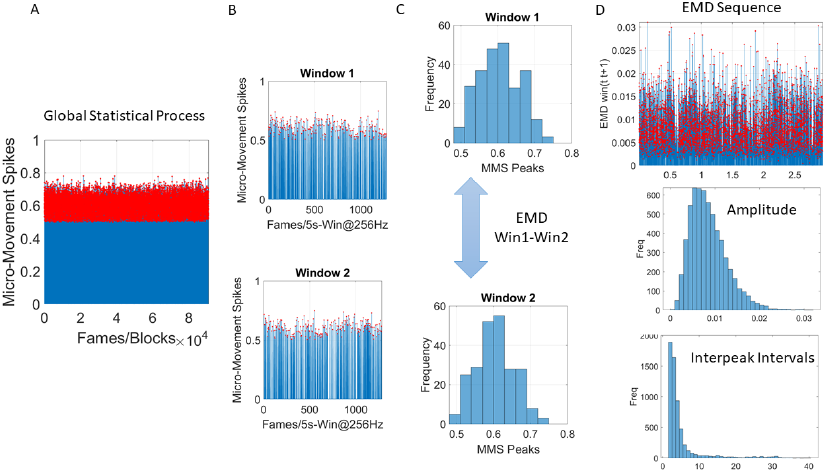
Global analyses by pooling the MMS across trials and blocks (A) and taking 5-second-long windows with 50% overlap (B) to obtain frequency histograms that can be compared using the EMD metric (C). (D) Sweeping through the full trajectory of a condition gives the EMD sequence to obtain the peaks in red and gather them into a frequency histogram tracking the fluctuations in amplitude of the EMD variation (i.e., how the distribution change shape and dispersion) and the rate at which these changes occur as the inter peak interval intervals measuring the distances as well across peaks representing the PDF transitions. These histograms are used in MLE estimation of the distribution parameters best describing this global process.

## Contribution

All authors had equal and complementary contributions to the work.

AV ran the experiment, recorded the original EEG data and analyzed behavioral results. EBT developed the novel analytical tools for EEG data that led to the new exploratory learning mode. Both authors contributed to the conceptualization of this work, interpretation of the results and the preparation of the manuscript. This work merges the motor control dynamic and stochastic perspective EBT, with the study of statistical learning AV.

