## Supplementary Figures for "Rethinking Statistical Learning as a Dynamic Stochastic Process, from The Motor Systems Perspective" (from another version of this preprint: version 1, 2022-01-19)

*Equal complementary contributions

^‡^ Corresponding Author

Additional Results on Global Analyses of PDF transitions evaluated through the EMD (as explained in Figure 11 of the Methods section of the main text.

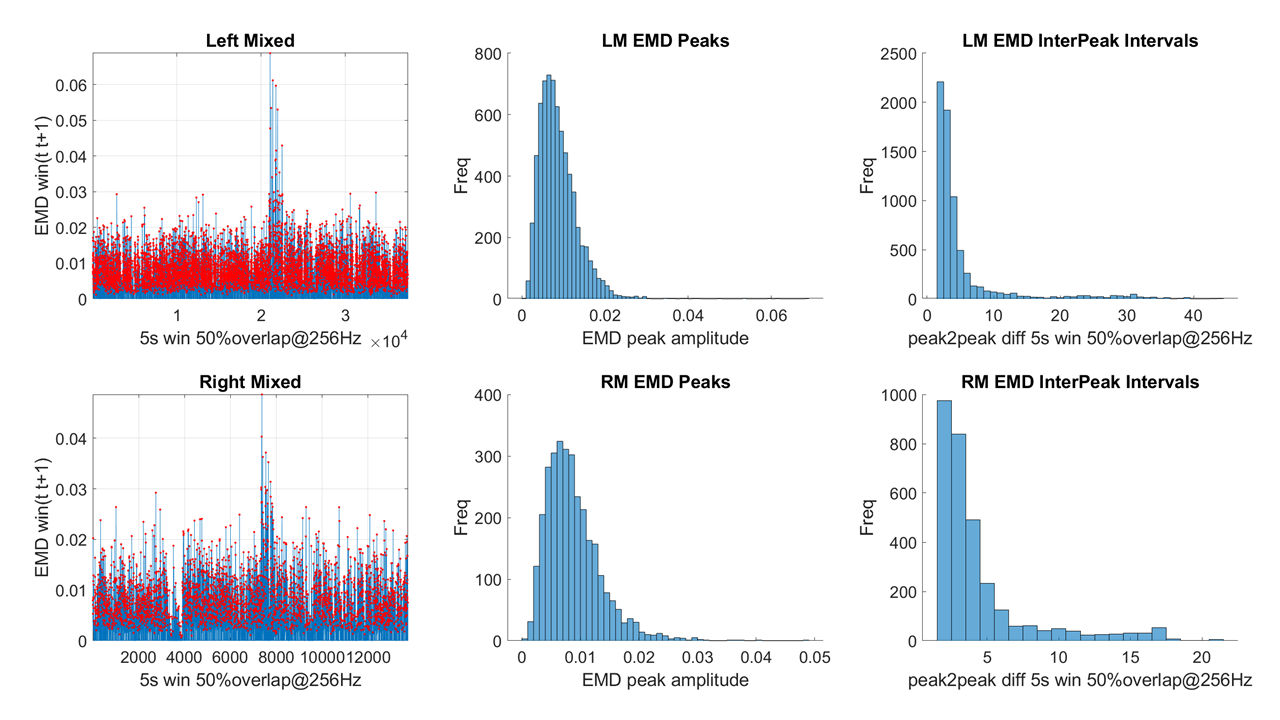

Supplementary Figure 1. Global analyses of the pooled data from all trials and blocks across participants for the mixed condition. Left and right oriented target trials on the top and bottom panels of the first column, respectively, showing the sequence of EMD values obtained by pooling across all data (order does not matter). Peaks (red) are used to build the frequency histograms depicted on the top (left target) and bottom (right target) panels of the second column. This are the fluctuations in EMD peak amplitude, quantifying the shifts in PDF empirically approximating the histograms using MLE. Panels on the third column show the frequency histograms of the interpeak intervals measuring the peaks’ distances from EMD window t and t+1, t represents the 5second-long window sweeping through the data with 50% overlap.

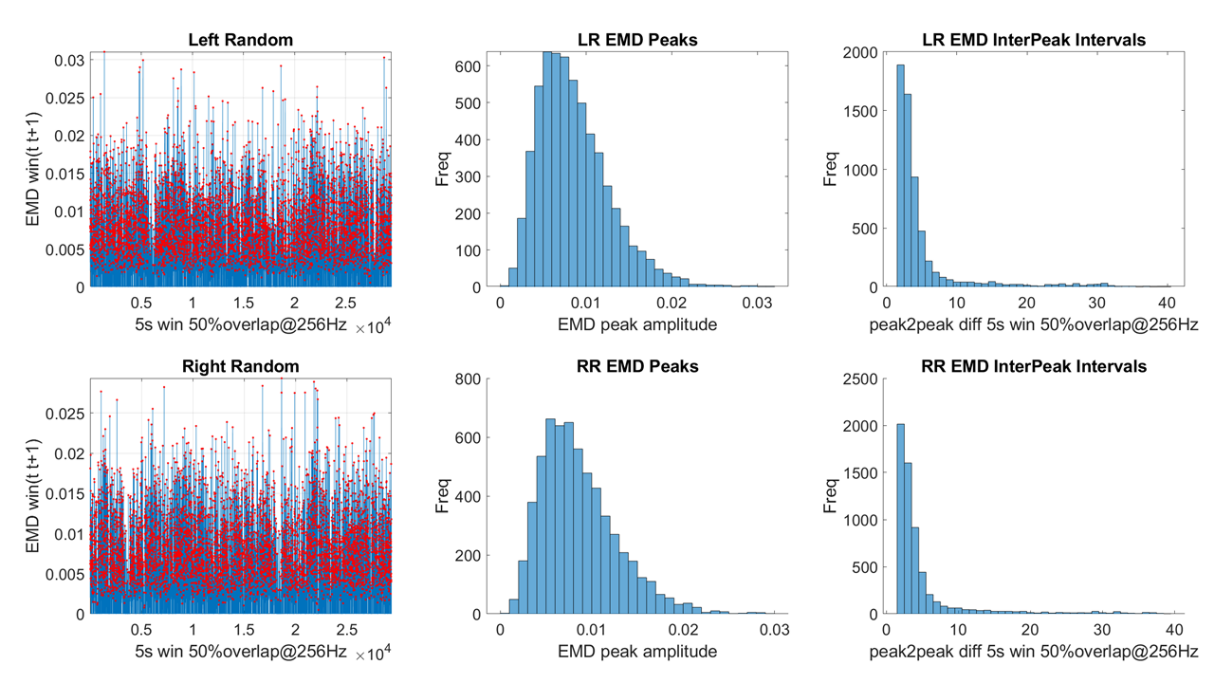

Supplementary Figure 2. Same as Supplementary Figure 1 for the random condition.

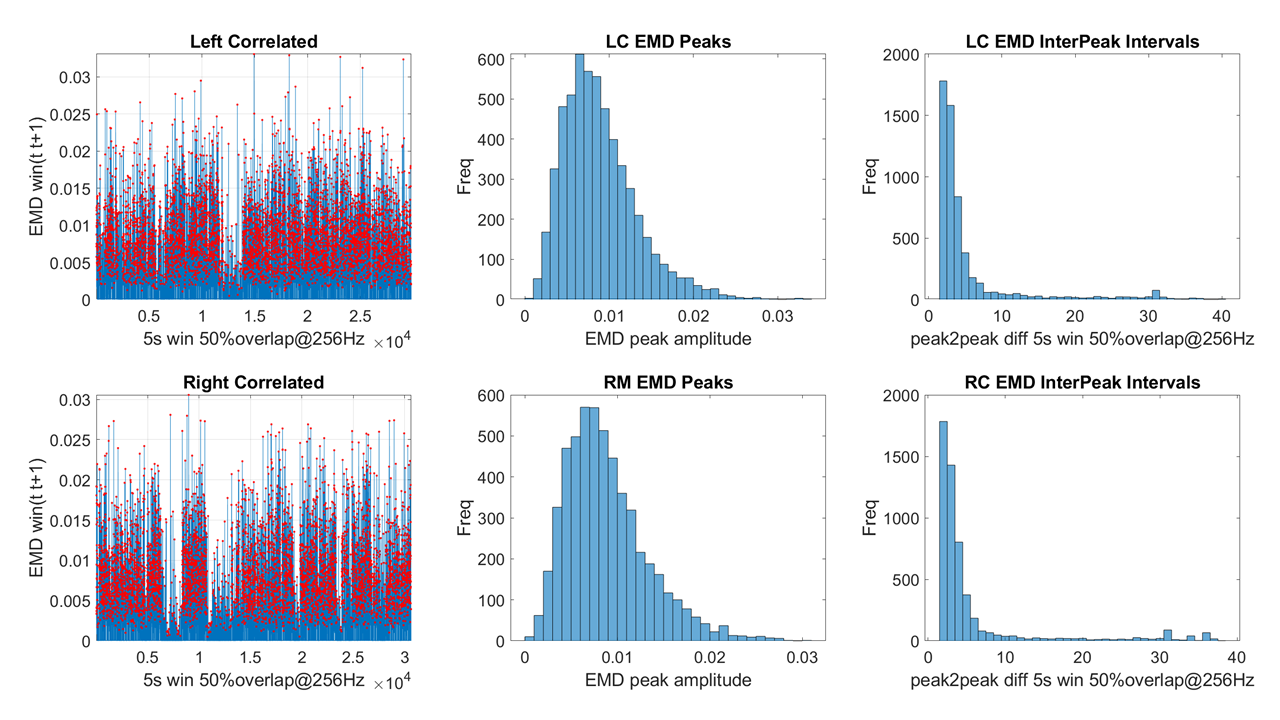

Supplementary Figure 3. Same as Supplementary Figure 2 for the correlated condition.

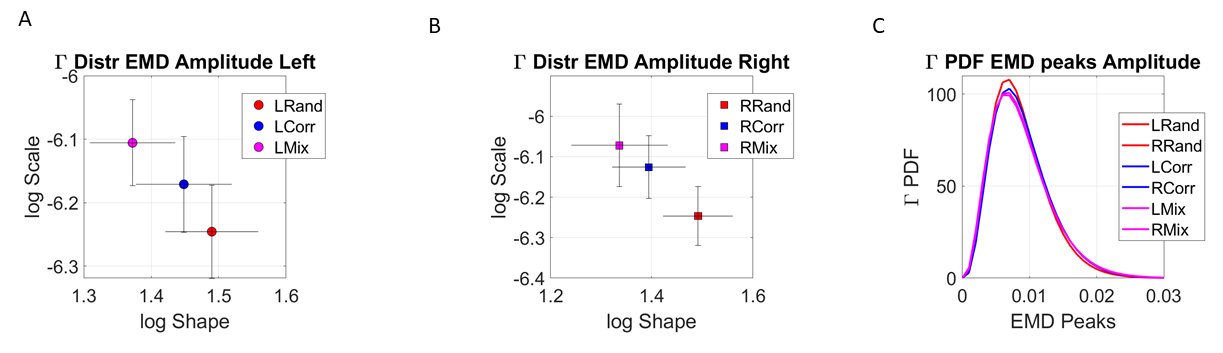

Supplementary Figure 4. Gamma process approximation of the global data for the fluctuations in amplitude. (A) Left oriented target global stochastic signatures mapped as empirically estimated parameters of the continuous Gamma family of probability distribution (see methods) plotted as points on the (shape, scale) parameter plane with 95% confidence intervals. (B) Same as in (A) for the right oriented target. (C) Corresponding PDFs estimated using MLE.

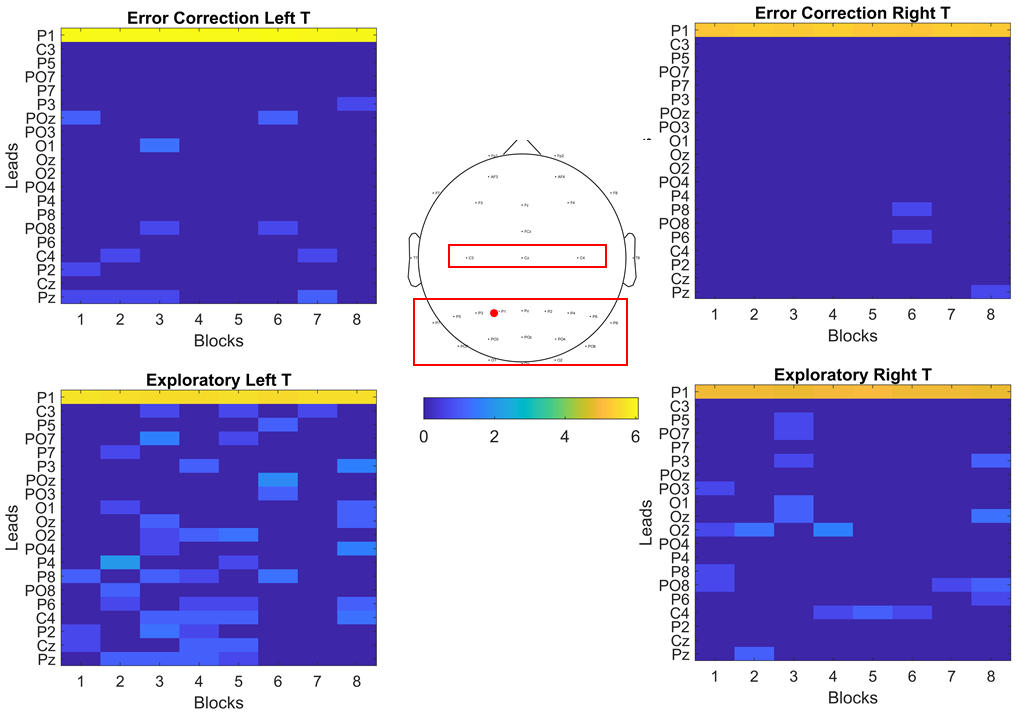

Supplementary Figure 5. Differentiation between error corrective and exploratory modes in the hubs’ activities across the leads under consideration. Center figure shows the leads used in the analyses. The leads acting as hubs of the network were quantified across trials and blocks (color bar in log scale) for each subgroup of participants, by pooling across participant type for all trials of each block in the mixed condition. The exploratory subgroup reveals a far more distributed code than the error corrective subgroup, with different patterns between target type. Notice as well the dynamic nature of this process, shifting from blocks 1 to 8.
